# Factors affecting pathogenicity of the turfgrass dollar spot pathogen in natural and model hosts

**DOI:** 10.1101/630582

**Authors:** R.A. Rioux, C.M. Stephens, J.P. Kerns

**Author notes:** Corresponding author (CMS). RR was the primary author. She designed, performed, and analyzed data for all experiments, and wrote the manuscript. CMS and JPK contributed ideas for experimentation and experimental design.

## Abstract

*Clarireedia* sp. (formerly called *Sclerotinia homoeocarpa*), the fungal pathogen that causes dollar spot of turfgrasses, produces oxalic acid but the role of this toxin in *Clarireedia* sp. pathogenesis is unknown. In the current study, whole plant inoculation assays were used to evaluate pathogenesis of *Clarireedia* sp. in various model hosts and investigate the role of oxalic acid in dollar spot disease. These assays revealed that both host endogenous oxalate content and pathogen-produced oxalic acid influence the timing and magnitude of symptom development. In time-course expression analysis, oxalate oxidase and related defense-associated germin-like protein genes in creeping bentgrass showed strong up-regulation starting at 48-72 hpi, indicating that germin-like protein genes are most likely involved in defense following initial contact with the pathogen and demonstrating the importance of oxalic acid in *Clarireedia* sp. pathogenesis. Overall, the results of these studies suggest that oxalic acid and host endogenous oxalate content are important for pathogenesis by *Clarireedia* sp. and may be associated with the transition from biotrophy to necrotrophy during host infection.

## Introduction

*Clarireedia* sp. (formerly called *Sclerotinia homoeocarpa*) cause dollar spot in turfgrass, which is one of the most important diseases of amenity turfgrasses worldwide [1]. Despite the previous name *Sclerotinia homoeocarpa*, recent literature confirmed that this pathogen is not a species of *Sclerotinia* or a member of any known genus within the *Rutstroemiaceae* [2]. Renaming and correct taxonomic placement of *Sclerotinia homoeocarpa* has resulted in the establishment of a new genus described as *Clarireedia* gen. nov. [2]. A total of four distinct pathogenic species that cause dollar spot have been characterized within this genus, with *C. jacksonii* and *C. monteithiana* primarily infecting C3 and C4 grasses in North America, respectively [2]. Isolates evaluated in the current study were sampled from both C3 and C4 grasses. Therefore, the *C. jacksonii/C. monteithiana* are referred to as *Clarireedia* sp. in this text. *Clarireedia* sp. have a broad host range that spans five plant families [2] and includes economically and ecologically important plants, such as switchgrass [3], perennial peanut [4], and tufted bulrush [5].

Dollar spot is of particular concern in golf course settings because the sunken, silver dollar-sized spots of diseased turf from which this disease earns its name are aesthetically unappealing and may negatively affect ball roll. Conditions favoring dollar spot development are broad. The fungus can grow and cause disease at temperatures ranging from 14-35°C, if relative humidity remains above 70% [1]. As a result, successful dollar spot management requires periodic fungicide applications throughout the growing season, making dollar spot the most economically important disease of turf worldwide [1, 6, 7].

Popular turfgrass species that are prone to dollar spot, particularly creeping bentgrass (*Agrostis stolonifera*, L.), have complex polyploid genomes and are outcrossing which make genetic characterization of dollar spot resistance difficult [8]. To date, only a single study exists comparing global gene expression among inoculated and non-inoculated creeping bentgrass plants [9] and dollar spot resistance has been identified on only a few major quantitative trait loci [10]. Moreover, the size of individual turfgrass plants makes it difficult to manipulate and visually assess disease phenotypes. An acceptable model host would provide a much-needed tool for molecular and phenotypic characterization of *Clarireedia* sp./host interactions and may be applicable to other turfgrass pathogens.

The vast array of genetic resources for *Arabidopsis thaliana* make it a natural choice as a model host for any pathosystem. Further, *A. thaliana* has been successfully used in a number of studies on pathogenicity of *Sclerotinia sclerotiorum* (*Ss*) and *Botrytis cinerea*, both of which belong to the same fungal order as *Clarireedia* sp. [11, 12, 13, 14, 15, 16]. However, *S. sclerotiorum* and *B. cinerea* are predominantly pathogens of dicots while *Clarireedia* sp. prefer monocot hosts, indicating that a monocot model host may be more suitable for studies of *Clarireedia* sp. pathogenesis. Rice is a forerunner in cereal genetic resources [17], but less closely related to creeping bentgrass and other major turfgrass species than *Brachypodium distachyon*, barley, or wheat [18, 19]. Studies of host/pathogen interactions and generous genetic resources are available for all three of these plants, indicating their potential utility as model hosts for *Clarireedia* sp. Consequently, one of the objectives of the present research was to compare *Clarireedia* sp. pathogenesis between these five model hosts and the natural host creeping bentgrass.

Identifying pathogenicity mechanisms is important for understanding pathogenesis in *Clarireedia* sp. Previously, researchers showed that this pathogen produces oxalic acid (OA) [12]. OA is a common phytotoxin and is produced by many important plant pathogens, particularly those in the *Sclerotiniaceae* family [20], to which *Clarireedia* sp. are closely related. *S. sclerotiorum*, a polyphagous necrotrophic plant pathogen with more than 400 hosts [6, 7] relies heavily on production of OA for successful pathogenesis. While the role of OA in *Ss* pathogenesis has been extensively studied [21, 15, 14, 7, 22], the function of OA in *Clarireedia* sp. pathogenesis is not clear.

Differences among the *Clarireedia* sp. and *S. sclerotiorum* host ranges indicate that these pathogens use OA in disparate ways during host colonization. *Clarireedia* sp. primarily infect monocot species within the family *Poaceae* [2]. Conversely, *S. sclerotiorum* infects a broad range of dicot species, but is not considered a pathogen on most monocots. Many grass species have oxalate oxidases that degrade OA, preventing successful infection by *Ss* [23, 24]. Transformation of dicot hosts with grass oxalate oxidase genes confers partial resistance to *Ss* [26, 27, 28]. Consequently, if OA is an important pathogenicity factor for *Clarireedia* sp., this fungus must use the phytotoxin in a manner that allows it to circumvent or overcome oxalate oxidase-mediated plant defenses. Both oxalate oxidase activity [9] and up-regulation of oxalate oxidase encoding genes occur during *Clarireedia* sp. infection of creeping bentgrass [29], indicating that OA is produced *in planta* during host colonization. A better understanding of the role of OA in host colonization by *Clarireedia* sp. and host oxalate oxidase-mediated defenses against *Clarireedia* sp. may contribute to novel disease management strategies.

Based on these observations, the objectives of the present research were to: 1) Investigate the infection process of *Clarireedia* sp. in creeping bentgrass and various model host systems and 2) determine the role of OA in *Clarireedia* sp./host pathogenesis.

## Materials and methods

### Biological materials

#### Plant materials

Monocots (S1 Table) were grown from seed and maintained in a growth room with a 14 h light period at 25±2°C and 10 h dark period at 22±2°C. *Arabidopsis thaliana* and *Nicotiana benthamiana* were grown under similar conditions, except growth chamber conditions were modified to an 18 h day-length and constant temperature of 26±2°C.

#### Fungal materials

Fungal isolates (S2 Table) were maintained on potato dextrose agar (PDA) under ambient temperature (22±2°C) in growth chambers with a 24 h dark period and transferred weekly to maintain viability. For long term isolate storage, mycelia were allowed to colonize sterile filter paper disks on PDA, air dried, placed in glass vials, and stored at −80°C.

### Inoculation assays and disease rating

Monocots were inoculated using the previously described parafilm sachet method [30]. The exact method used differed slightly with host species (S1 Fig). Dicots were inoculated by placing an agar plug mycelia side down on a fully expanded leaf surface. Flats containing inoculated dicots were covered with plastic domes to promote relative humidity >90%. Parafilm alone maintained high relative humidity for monocot inoculations and domes were not needed. Four-day-old cultures of *Clarireedia* sp. were used for all plant inoculations. Control plants were mock-inoculated with agar plugs from fresh PDA plates.

Symptom severity was rated every 24 h using a rating scale modified from the Horsfall-Barratt scale (S3 Table) [31]. This method allowed for direct comparison of symptom severity between species with diverse physical characteristics and symptom phenotypes. To prevent the possibility of rater-to-rater variability all symptom severity ratings were made by the same individual.

Inoculation experiments were repeated three times with three replicates of each treatment or treatment combination per experimental repetition (n=9). Data were analyzed with experimental repetition treated as a random blocking factor and blocks within each repetition treated as a random factor nested within experimental repetition. Plant species and fungal isolate were both considered fixed effects.

#### Trypan blue staining and microscopy

Inoculation and sample collection for all plant hosts and *Clarireedia* sp. isolates were performed in parallel. Scissors were used to excise plant material surrounding the site of inoculation and individual tissue samples were placed in 2mL microcentrifuge tubes. Staining and clearing of tissues were performed as previously described [32]. All samples were visualized with an Axio Scope.A1 compound microscope (Zeiss, Thornwood, NY). Images were captured with an AxioCam MRc camera and processed with accompanying AxioVision Real 4.7.1 imaging software (Zeiss, Thornwood, NY). A minimum of three samples were examined for each time-point, isolate, and host combination.

#### Whole plant oxalate quantification

Plants for oxalate quantification were inoculated as previously described. For fresh plant samples, 0.25 g of tissue were ground in liquid nitrogen with a mortar and pestle then added to a 15 mL centrifuge tube containing 4 mL of 0.2 M KH_2_PO_4_ buffer (pH 6.5) [33]. Tubes containing ground plant tissue were placed on their sides on an incubator shaker table and mixed overnight with constant gentle shaking at 90 revolutions per minute at room temperature (22±2°C). The following day, 2 mL of the liquid portion from each sample tube was transferred to a fresh 15 mL centrifuge tube and mixed with 2 mL of freshly prepared oxalate quantification kit sample diluent (Trinity Biotech, Jamestown, NY). The remainder of the oxalate assays were performed according to the manufacturer’s instructions (Trinity Biotech, Jamestown, NY). Oxalate content was quantified based on absorption at 590 nm with a UV/Vis spectrophotometer (Beckman Coulter, Brea, CA). A 0.5 mmol oxalate standard was included in each run.

Due to the high volume of samples in time-course oxalate studies, samples for this assay were harvested, weighed, flash frozen in liquid nitrogen, and stored at −80°C until use. Pilot studies indicated that flash freezing did not affect oxalate content in comparison with samples processed when fresh. Frozen samples were lightly ground in liquid nitrogen with a mortar and pestle, then transferred to Lysing Matrix A FastPrep tubes (MP Biomedicals, Santa Ana, CA) with 1.5 mL KH_2_PO_4_ buffer and ground in a FastPrep at 6.0 m/s for 40 s (MP Biomedicals, Santa Ana, CA). Homogenized samples were poured into 15 mL centrifuge tubes and mixed with 1.5 mL sample diluent. Oxalate quantification was then performed as previously described.

### Germin-like protein gene expression analysis

Pots of one-week-old creeping bentgrass turfgrass cultivars ‘96-2’ and ‘Focus’ were inoculated with two agars plugs colonized with *Clarireedia* sp. mycelia and covered to maintain humidity >90%. Cultivars ‘96-2’ and ‘Focus’ were chosen for their low and high resistance to dollar spot in field trials, respectively (http://www.ntep.org). The experiment was arranged as a two treatment (inoculated and mock-inoculated) by two cultivar (’96-2’ and ‘Focus’) factorial within a randomized complete block design with four replications. Pots were resampled at 0, 24, 48, 72, and 96 hpi by harvesting 100 mg of tissue from the area surrounding the site of inoculation, removing roots and leaf blade tips, flash freezing in liquid nitrogen, and storing at −80°C until use.

Samples for RNA extraction were ground with a mortar and pestle in 500 µL frozen Trizol reagent then transferred to 2 mL microcentrifuge tubes containing 500 µL Trizol reagent. RNA extraction was then performed with the Trizol Plus RNA Purification Kit including the optional on-column DNase step according to the manufacturer’s instructions (Life Technologies, Grand Island, NY). RNA quality was assessed with an Experion Automated Electrophoresis Station using the Experion RNA StdSens analysis kit (BioRad, Hercules, CA) and yield was quantified with an ND-1000 spectrophotometer (Nanodrop, Wilmington, DE). Samples were stored at −80°C until further use.

Synthesis of cDNA and RT-qPCR were performed according to the specifications of the MIQE guidelines. Prior to cDNA synthesis, a no cDNA control RT-qPCR run was performed on pure RNA to confirm the absence of contaminating genomic DNA. RNA samples confirmed as free of genomic DNA were diluted to equivalent concentrations and subjected to cDNA synthesis with iScript cDNA Synthesis Kit according to the manufacturer’s instructions (BioRad, Hercules, CA). All cDNA synthesis reactions for a single time-point were performed simultaneously to decrease any inherent variability in the synthesis process. cDNA quality was assessed by comparing Cq values for potential reference genes and reference genes were selected using BestKeeper Software [34]. Creeping bentgrass actin (ACT7) and glyceraldehyde phosphate dehydrogenase (GAPDH) genes were selected as stable reference genes for all time-points in both cultivars.

Primers for RT-qPCR were designed from publicly available creeping bentgrass EST sequences with Primer3 primer design software (http://www.genome.wi.mit.edu/genome_software/other/primer3.html; S4 Table) and validated by generating efficiency curves from a dilution series of cDNA pooled from all time-points. For RT-qPCR, cDNA samples were diluted 10-fold and a final volume of 8 µL cDNA was used in each of three 20 µL technical replicates per reaction, along with 10 µL SsoFast EvaGreen Supermix (BioRad, Hercules, CA), 0.3 µM of each primer, and nuclease free water. All reactions were run in hard-shell skirted 96-well plates with a white top and clear wells and sealed with Microseal ‘B’ adhesive film (BioRad, Hercules, CA). A CFX-96 detection system (BioRad, Hercules, CA) was used for quantification. PCR conditions were 98°C for 2 m; 40 cycles of 98°C for 2 s and 55°C for 5 s; followed by a dissociation curve with 80 cycles starting at 70°C for 10 s and increasing in 0.2°C increments per cycle. The dissociation curve was used to detect primer-dimers and amplification of a single product within the CFX detection system software (BioRad, Hercules, CA).

To calculate relative expression of target genes, transcript abundance was internally normalized to the two reference genes (actin and elongation factor 1-α) that were included on each experimental plate using the formula 2^Δ*Cq*(target-reference)^ [35]. Relative expression ratios of target genes were then calculated using a method based on the previously described 2^−ΔΔ*Cq*^ method [35, 36, 37]. Briefly, normalized target gene Cq values for infected plant tissue were divided by a calibrator value, which was calculated by averaging the normalized Cq values for all uninfected 96-2 samples from the same time point [35]. This method allowed for comparison between all treatment levels at each time-point. Statistical analysis was performed on resulting relative expression ratios using the proc mixed procedure in SAS v.9.3 (SAS Institute, Cary, NC). Orthogonal contrasts were used to compare relative expression ratios between mock-inoculated and inoculated samples within each cultivar and between the inoculated samples of the two cultivars.

### Statistical analysis

Data analysis was performed using the MIXED, generalized linear mixed model (GLIMMIX), and regression (REG) procedures in SAS version 9.3. Initial models were fit including all random factors and interactions with fixed factors. Model fitting criteria (Akaike’s Information Criterion) were then used to remove random factors and interactions not contributing to variability in order to identify the best model for each data set. Visual analysis of residual plots and output from PROC UNIVARIATE were used to assess normality of the data. Data were transformed as necessary to best approximate the normal distribution using the DIST= option in PROC GLIMMIX. Reported values were back-transformed using the ILINK option. The ANOVA F-test was used to determine contributions of fixed factors to variability in the measured response variable(s). Additionally, pre-planned orthogonal contrasts were used to compare between specific fixed factors.

Use of the symptom severity rating scale resulted in data with a non-normal distribution that could not be corrected through common data transformations available within PROC GLIMMIX. Consequently, a Friedman’s two-way analysis for block designs was employed [38]. For each experimental repetition, the symptom severity for each data point (species by treatment) was ranked within blocks previously defined during experimental set-up. The resulting ranks were assessed for normality and agreement between experimental repetitions. The rank data from each experiment repetition was then pooled and analysis of variance was performed with the rank output as the response variable and both repetition and block(repetition) as random factors.

## Results

### Characterization of the *Clarireedia* sp. infection process on natural and model hosts

Microscopic characterization of infection at various time-points following inoculation was similar regardless of host (Fig 1). As early as six hours-post inoculation, *Clarireedia* sp. hyphae were observed growing along cell walls (Fig 1A) and initiating host penetration through infection pegs (Fig 1A) and stomata (Fig 1B). This resulted in extensive colonization of host tissue, particularly xylem, shortly following inoculation (Fig 1C-E). Host cell penetration and colonization continued for at least 72 h. Dead host cells, as indicated by trypan blue staining, were observed at minimum of 48 hrs after inoculation (Fig 1J-K). Differences in the infection process were not observed with the four *Clarireedia* sp. isolates used in this study.

**Fig 1.**
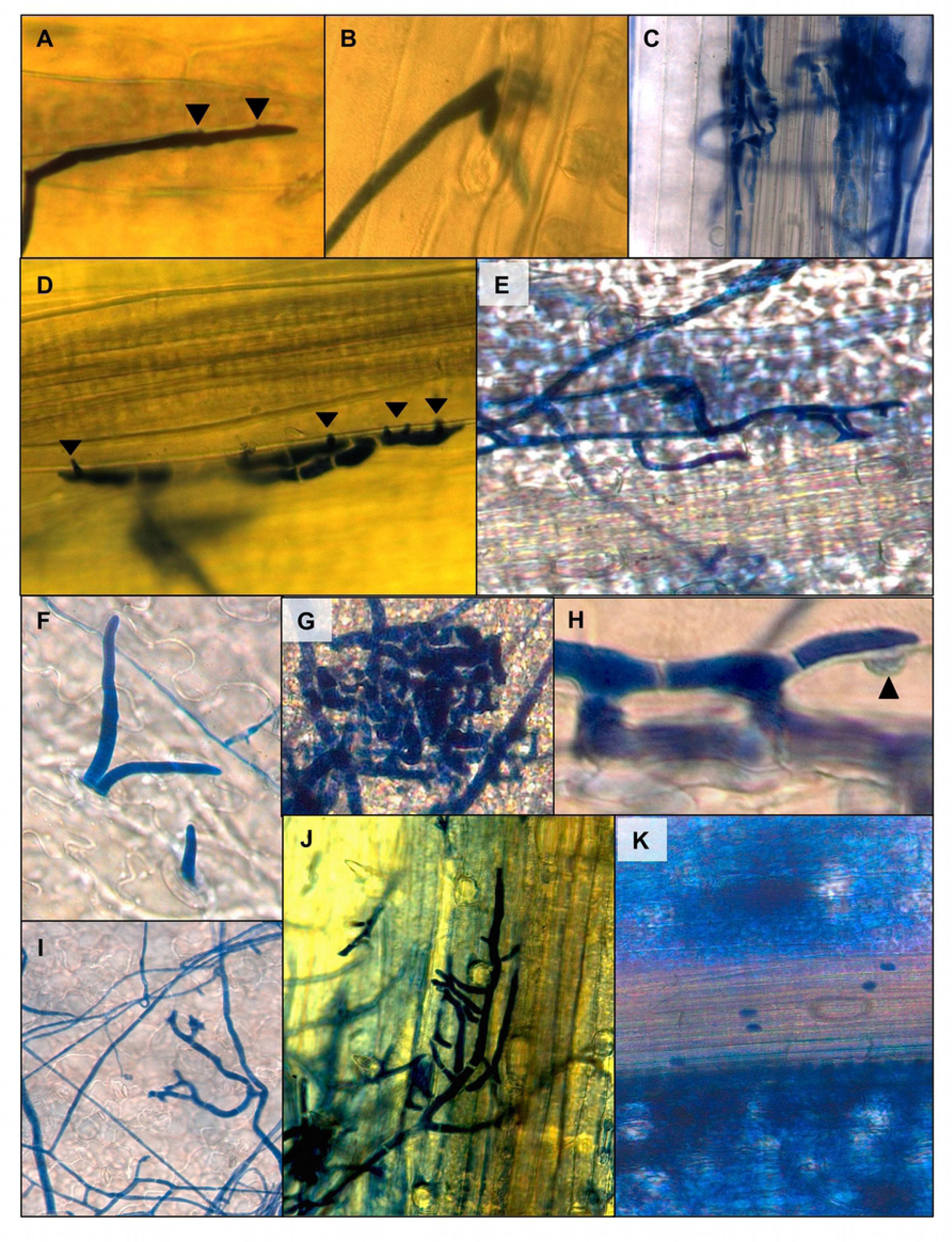
Microscopic analysis of plant foliar tissue colonization by *Clarireedia* sp. In-tact plants were inoculated and plant samples were taken by excising the area around the site of inoculation. All samples were cleared of chlorophyll and stained with 0.01% trypan blue. Host penetration was observed as early as 6 hpi and included: **A)** Growth along cell walls and formation of infection pegs (arrowheads) projecting directly into host tissue; shown in creeping bentgrass. **B)** Direct penetration through stomata; shown in barley and **C)** Extensive colonization of host tissue with an affinity for vascular tissue; shown in barley. At 12 to 24 hpi fungal hyphae continued to grow along cell walls and form penetration pegs into vascular tissue, as observed in **D)** Creeping bentgrass at 12 hpi **E)** Wheat at 24 hpi and **G)** Rice at 24 hpi. **F)** *Clarireedia* sp. protruding from a stomate in *Arabidopsis thaliana* tissue colonized by fungal mycelia. **H)** Appressorium (arrowhead) formed by a *Clarireedia* sp. hypha on barley at 96 hpi. **I)** Extensive host cell death, as indicated by trypan blue staining outside of fungal hyphae, was not observed until 4-7 dpi regardless of host. **J)** Extensive cell death 7 dpi inoculation of *Brachypodium distachyon* with *Clarireedia* sp. **K)** Extensive cell death outside of host vascular tissue at 7 dpi in wheat. Images are representative of similar observations made for all hosts and *Clarireedia sp.* isolates used in this study.

Symptoms of *Clarireedia* sp. infection were similar in monocot hosts, but differed in the dicot host *Arabidopsis thaliana* (Fig 2). Inoculation of creeping bentgrass plants resulted in bleaching and necrosis of the inoculated leaf area (Fig 2F). Within this area, individual leaf blades with hourglass shaped lesions of white-gray tissue surrounded by reddish-brown borders were identified (Fig 2E). Similar lesions were observed on the four model monocot hosts (Fig 2A-D). Occasionally, complete wilting of infected leaves was observed, particularly in the later stages of infection. Symptoms on *A. thaliana* began as water-soaked lesions, with the affected areas eventually wilting. These lesions were visually distinct from necrotic lesions typically observed in studies of *A. thaliana* inoculated with the related pathogen *Ss* [15]. The petiole of the inoculated *A. thaliana* leaf would become necrotic, constricted, and the symptomatic leaf would often separate from the plant following severe infection.

**Fig 2.**
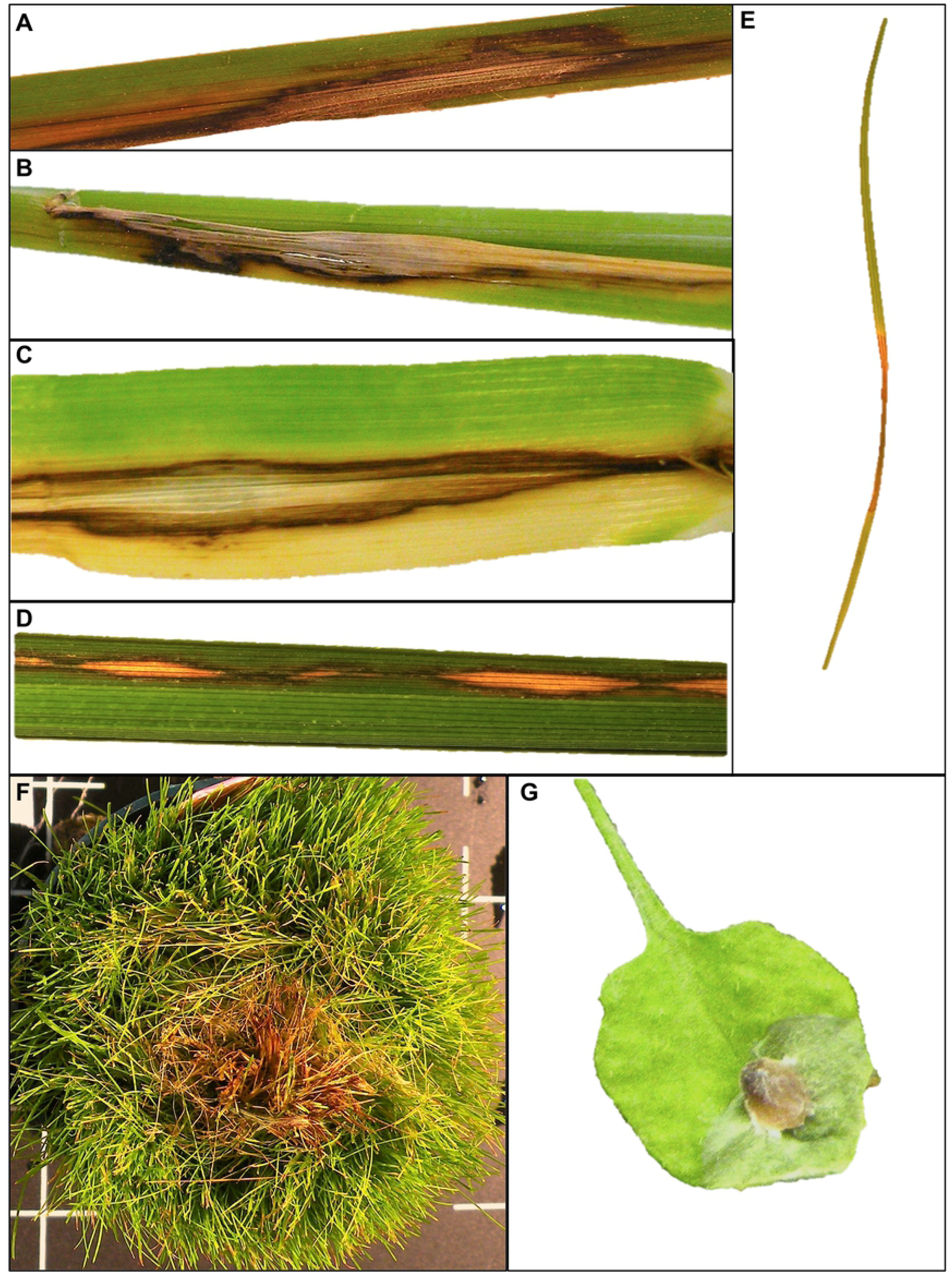
Symptoms produced by *Clarireedia* sp. on natural and model host plants. *Clarireedia* sp. infected and produced visible symptoms on all host plants. Similar symptoms were observed on **A)** *Brachypodium distachyon* **B)** Wheat **C)** Barley and **D)** Rice. These symptoms included light tan to white lesion with reddish brown borders and resembled those observed on **E)** Creeping bentgrass, a natural host. **F)** Creeping bentgrass stand symptoms resulting from our parafilm sachet inoculation method looked visibly similar to symptoms observed in the field. **G)** Foliar symptoms observed on *Arabidopsis thaliana* were distinct from those observed on monocot hosts and were characterized by water soaking and tissue collapse.

Analysis of area under the disease progress curve (AUDPC) values revealed a significant effect of species, but not isolate or isolate × species interaction based on a simple two-way ANOVA F-test *(P=*0.01) (Fig 3A). In particular, AUDPC values were different between rice and creeping bentgrass according to a Dunnett’s test comparing mean AUDPC values for species (*P*=0.0043). Overall symptom severity in time-course assays was only affected by host species at 48 hpi *(P=*0.03), when symptoms began to rapidly develop in *B. distachyon* and *O. sativa* (Fig. 3B). However, greater symptom severity was observed in rice at 48, 72, and 96 hpi (*P*<0.05) and in *B. distachyon* at 48 hpi (*P*=0.01) when compared specifically to creeping bentgrass. No additional differences in symptom severity between creeping bentgrass and other species were found. Throughout these experiments, symptom severity on mock-inoculated controls remained at or near zero.

**Fig 3.**
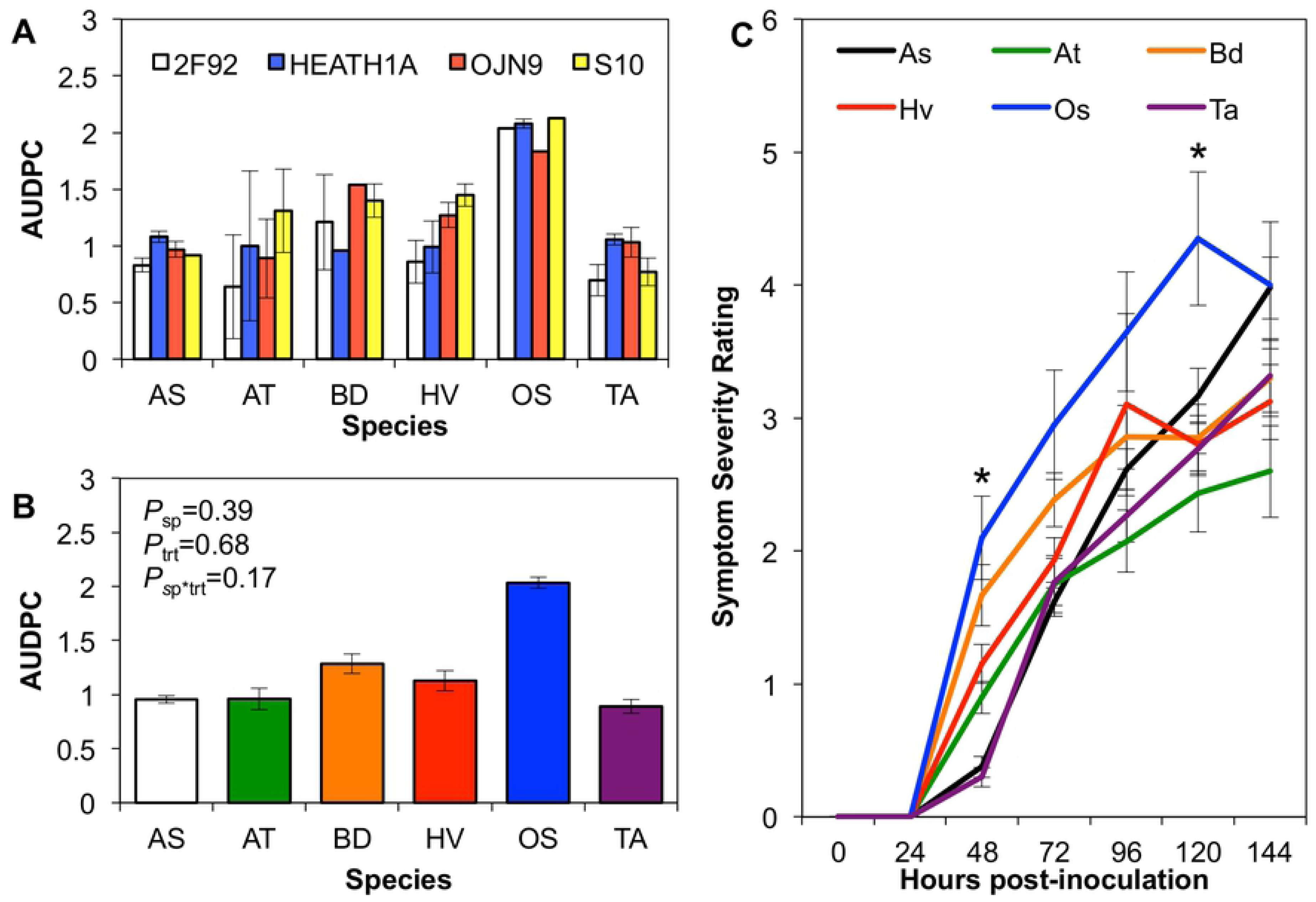
Symptom severity of *Clarireedia* sp. infection on natural and model hosts. All plants were inoculated in the same time in a split-plot RCBD with host as the whole-plot factor and *Clarireedia* sp. isolate as the sub-plot factor. Three experimental repetitions were conducted and treated as a random blocking factor in experimental analyses. Blocks within experimental repetitions were considered random factors nested within repetition. Host, isolate, and host × isolate interaction were fixed effects. **A)** AUDPC for each *Clarireedia* sp. isolate on the six host plants. Colored bars represent four different *Clarireedia* sp. isolates. P-values presented are from a two-way ANOVA with α = 0.05. Asterisks indicate significant difference from the natural host creeping bentgrass (AS) based on a Dunnett’s test with α = 0.05. Only the AUDPC for rice (OS) differed from that of creeping bentgrass. N≥9 for each species*isolate combination. **B)** Time-course progression of infection by *Clarireedia* sp. on all host species. The black asterisk indicates an overall significant effect of species on symptom severity at 48 hpi, as assessed by one-way ANOVA. Blue and orange asterisks indicate significant differences between rice and *B. distachyon* (BD), respectively, and creeping bentgrass at the corresponding time-points based on analysis with Dunnett’s test. N≥30 for each species. Abbreviations for both figures are as follows: AS=*Agrostis stolonifera* (creeping bentgrass); AT=*Arabidopsis thaliana*; BD=*Brachypodium distachyon*; HV=*Hordeum vulgare* (barley); OS=*Oryza sativa* (rice); and TA=*Triticum aestivum* (TA). For significance codes, *=P≤0.05; **=P≤0.01; ***=P≤0.001. All error bars represent ± one standard error of the mean.

### Relationship between host endogenous oxalate content and symptom development

To explore the relationship between *Clarireedia* sp. pathogenicity and OA, the oxalate content of plants inoculated with *Clarireedia* sp. or mock-inoculated with PDA was compared. A positive relationship between oxalate content and symptom severity was detected in creeping bentgrass, barley, and wheat, with R^2^ values ranging from 0.33 to 0.52 (Fig 4). The relationship between oxalate content and symptom severity in rice and *B. distachyon* was also positive, but lower than for the other species (R^2^ values of 0.10 and 0.01, respectively). A negative relationship between oxalate content and symptom severity was observed in *Arabidopsis thaliana*; however, correlation was low. Endogenous oxalate content of the six species differed significantly (*P*<0.0001) and comparisons against creeping bentgrass revealed that both rice and *B. distachyon* had higher oxalate levels (*P*<0.0001; Fig 4).

**Fig 4.**
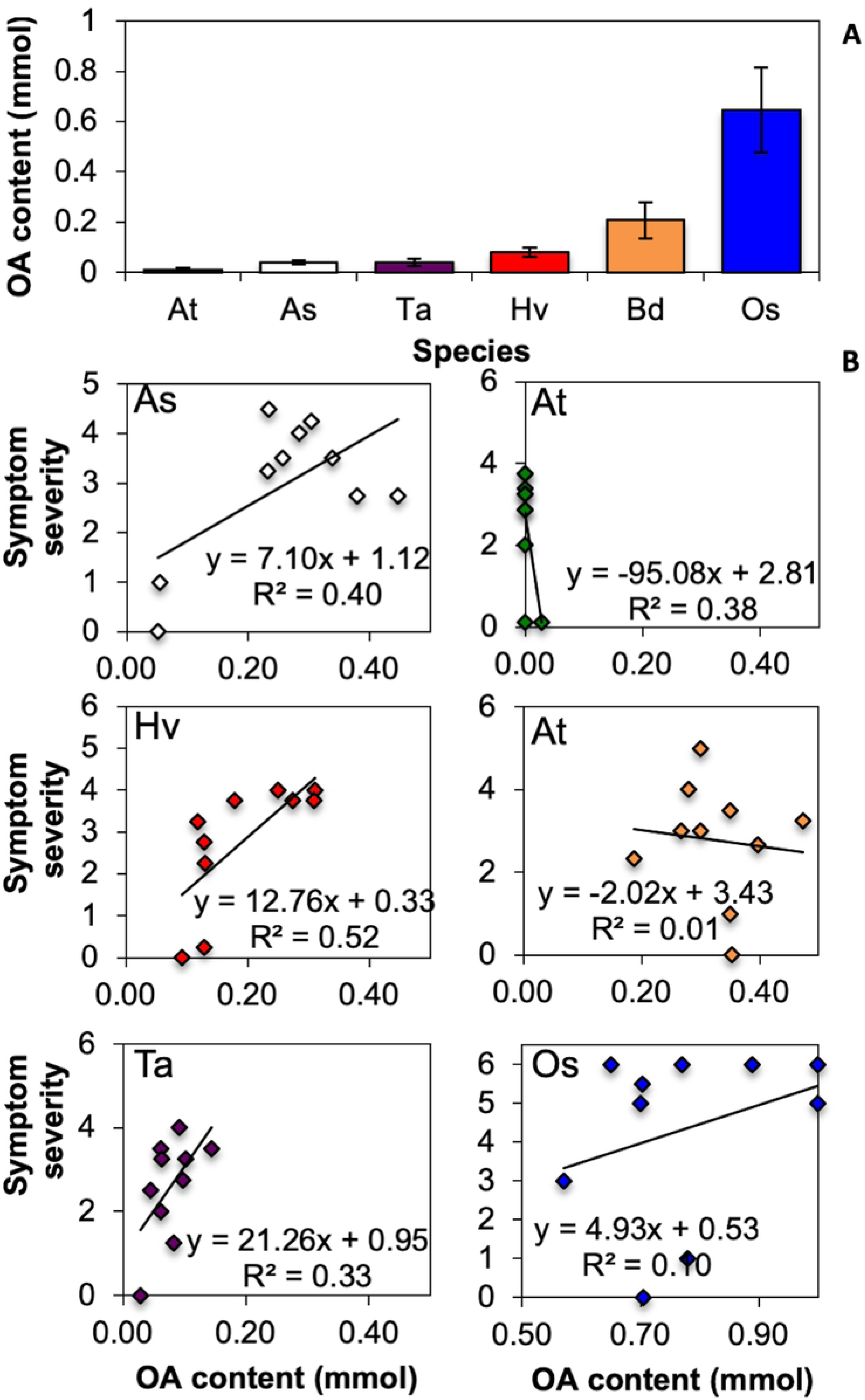
Correlation between host endogenous oxalate content and symptom severity. **A)** Endogenous oxalate content of all host species used in this study. Asterisks indicate significant difference from creeping bentgrass (AS) at the α=0.05 level using Dunnett’s test. Error bars represent ± one standard error of the mean. **B)** Correlation between oxalate content and symptom severity of *Clarireedia* sp. infection were fit a linear regression model for creeping bentgrass/*Agrostis stolonifera* (As), *Arabidopsis thaliana* (At), *Brachypodium distachyon* (Bd), barley/*Hordeum vulgare* (Hv), rice/*Oryza sativa* (Os), and wheat/*Triticum aestivum* (Ta). All regression equations, R^2^, and P-values were obtained from models fit with SAS proc reg. For significance codes, *=P≤0.05; **=P≤0.01; ***=P≤0.001. All error bars represent ± one standard error of the mean.

A time-course experiment was performed to compare symptom severity and oxalate content between *Clarireedia* sp.-inoculated and mock-inoculated plants during the progression of infection. Creeping bentgrass was selected to represent species with low oxalate content and *B. distachyon* was selected to represent species with high oxalate content. *B. distachyon* was selected over *O. sativa* because symptom severity and genetics were more similar to creeping bentgrass [18]. Oxalate content in *Clarireedia* sp.-inoculated plants gradually increased over time and paralleled the development of symptoms (Fig 5A). Oxalate content in mock-inoculated plants remained low throughout the experiment. Significant differences between the oxalate content in *Clarireedia* sp. and mock-inoculated creeping bentgrass developed at 48 hpi (*P*=0.004) and 120 hpi (*P*<0.0001). Conversely, oxalate content of inoculated and mock-inoculated *B. distachyon* plants remained relatively stable over time (Fig 5C). The oxalate content of inoculated and mock-inoculated *B. distachyon* only differed significantly at 120 hpi (p=0.003). Oxalate content correlated to symptom severity in both hosts (*P<0.0001*), but 75% of the variability in symptom severity was explained by oxalate content in creeping bentgrass, compared to only 28% in *B. distachyon*.

**Fig 5.**
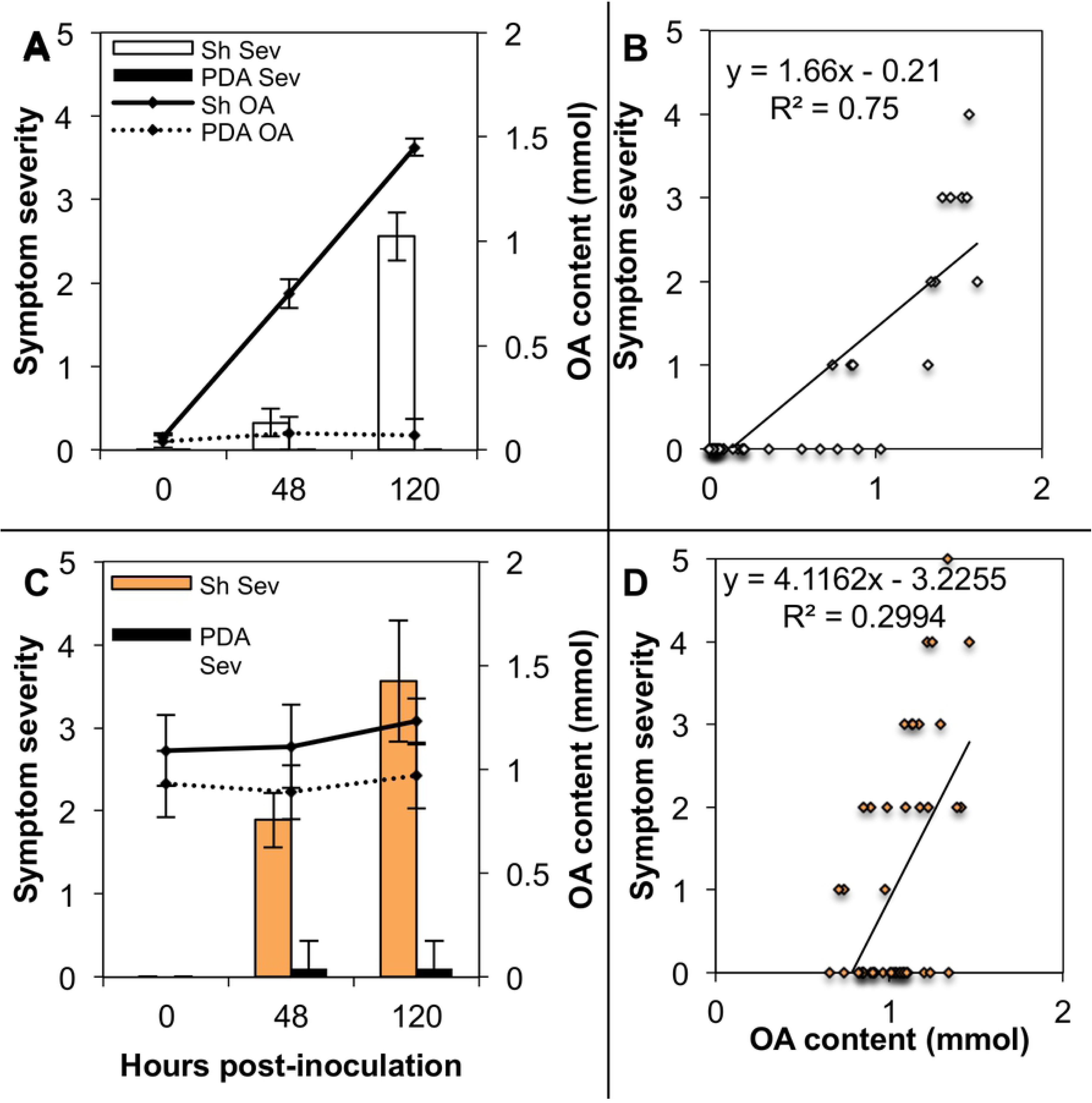
Time-course for the relationship between symptom severity and oxalate content in creeping bentgrass and *Brachypodium distachyon.* **A)** Time-course of symptom severity and oxalate content in creeping bentgrass. Asterisks indicate difference between oxalate content in *Clarireedia* sp. and mock-inoculated plants by one-way ANOVA with a cut-off value of α=0.05. Errors bars represent ± one standard error of the mean. **B)** Scatter plot showing the correlation between oxalate content and symptom severity for creeping bentgrass. Regression line calculated from simple linear regression in SAS proc reg. **C)** Time-course of symptom severity and oxalate content in *B. distachyon*. Asterisks indicate difference between oxalate content in *Clarireedia* sp. and mock-inoculated plants by one-way ANOVA with a cut-off value of α=0.05. Errors bars represent ± one standard error of the mean. **D)** Scatter plot showing the correlation between oxalate content and symptom severity for *B. distachyon*. Regression line calculated from simple linear regression in SAS proc reg. For significance codes, *=P≤0.05; **=P≤0.01; ***=P≤0.001. All error bars represent ± one standard error of the mean.

To further elucidate the influence of pathogen-produced OA relative to plant endogenous oxalate content on symptom development in creeping bentgrass and *B. distachyon*, the aggressiveness of *Clarireedia* sp. isolates producing varying levels of OA in culture was compared (Fig 6). Isolates A1421, HE10G19, and ML75 produced high, low, and moderate amounts of OA, respectively, based on acidification of bromophenol blue amended medium assays (Fig S2) [39]. Orthogonal contrasts were used to compare symptom severity between HE10G19 (low OA prod) and the remaining two isolates at each time point (Fig 6). On creeping bentgrass, aggressiveness of isolates differed at 72 hpi. From this point on, HE10G19 was less aggressive than the other two isolates (p≤0.05; Fig 6B). Aggressiveness among the *Clarireedia* sp. isolates did not vary on *B. distachyon* (Fig 6C).

**Fig 6.**
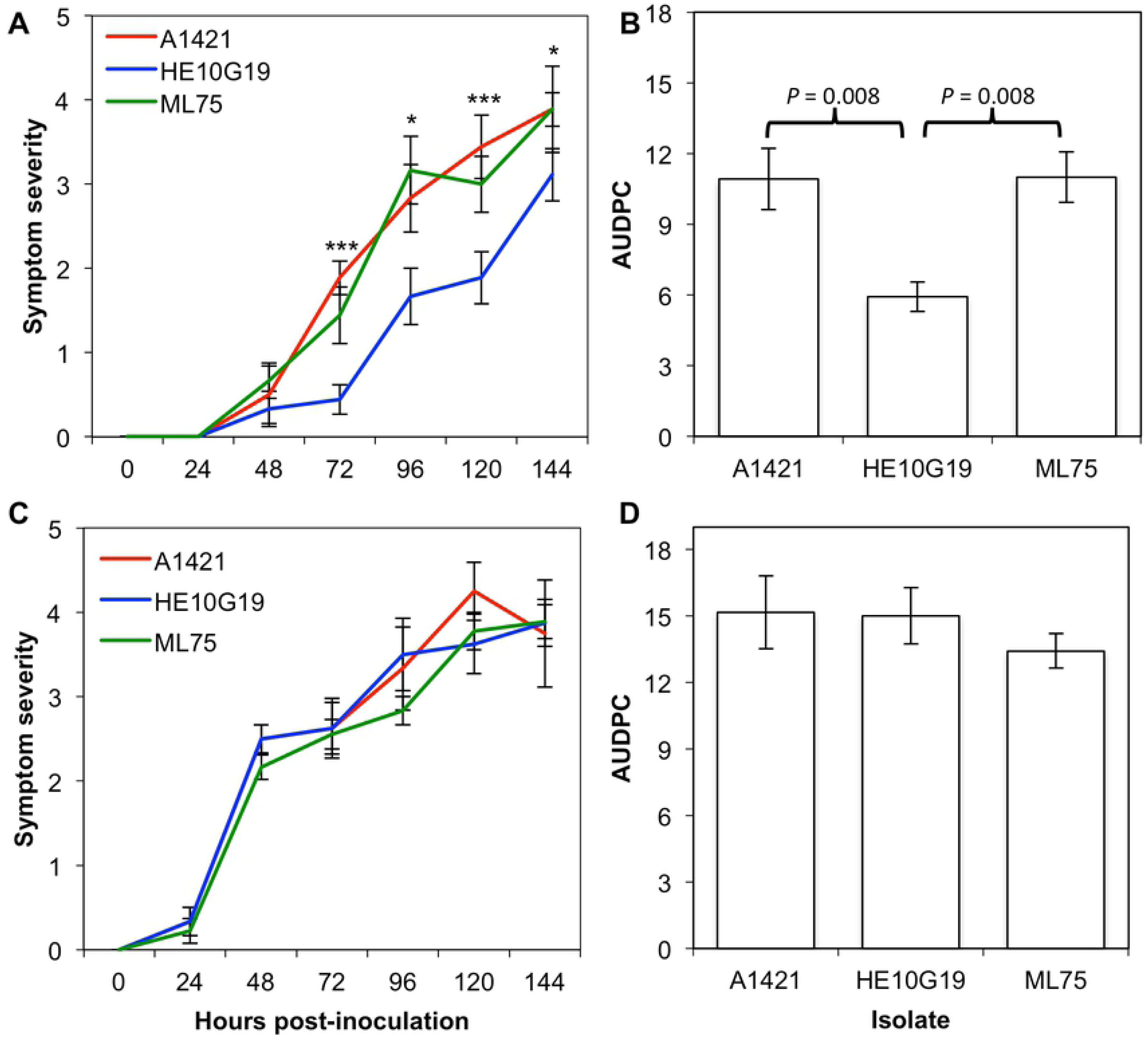
Progression of infection on creeping bentgrass and *Brachypodium distachyon* by *Sclerotinia homoeocarpa* isolates with varying oxalic acid production capacities. Isolates producing high, moderate, and low amounts of oxalic acid were selected based on their ability to produce a color change in pH indicator-amended media (S2 Fig). In line graphs, red, green, and blue lines indicate isolates with high, moderate, and low oxalic acid production abilities, respectively. Creeping bentgrass and *B. distachyon* experiments were performed in tandem. Values shown in these graphs represent the means from the three experimental repetitions. **A)** Time-course of symptom development and **B)** AUDPC of symptom severity on creeping bentgrass. **C)** Time-course of symptom development and **B** AUDPC of symptom severity on *B. distachyon.* Asterisks in **A** represent a significant effect of isolate in one-way ANOVA with α=0.05. P-values in **B** were obtained through orthogonal contrasts comparing the AUDPC means between the isolates indicated. No difference between isolates was detected for *B. distachyon*. For significance codes, *=P≤0.05; **=P≤0.01; ***=P≤0.001. All error bars represent ± one standard error of the mean.

### Comparison of oxalate oxidase expression in creeping bentgrass cultivars

A previous study determined various germin-like protein (GLP) genes, including oxalate oxidases, were some of the most up-regulated genes in the dollar spot susceptible cultivar ‘Crenshaw’ 96 hpi with *Clarireedia* sp. [29]. In the present study, RT-qPCR was used to compare the expression of oxalate oxidase and other germin-like protein genes between cultivars considered resistant and susceptible to *Clarireedia* sp. Four genes of interest were selected for RT-qPCR analysis of expression through sequence alignment and phylogenetic analysis of creeping bentgrass ESTs and known GLP genes from barley, rice, and *B. distachyon* (S3 Fig). Expression was tested by extracting total RNA from four independent pots of *Clarireedia* sp.*-*inoculated or mock-inoculated resistant and susceptible creeping bentgrass cultivars. All four of these genes were up-regulated in both resistant and susceptible cultivars by 96 hpi in *C. Clarireedia* sp.*-* inoculated versus mock-inoculated control plants (Figs 7 and S3). The oxalate oxidase gene, AST_798, was up-regulated in the susceptible cultivar at 48, 72, and 96 hpi. Expression of AST_798 was stronger in the susceptible cultivar when compared to the resistant cultivar 48 and 72 hpi. AST_798 was only up-regulated in the resistant cultivar at 96 hpi (Fig 7A). Similarly, AST_854, a creeping bentgrass gene grouping with defense-associated GLP genes from barley and rice in phylogenetic analysis, was up-regulated in the susceptible cultivar at 72 and 96 hpi, but only at 96 hpi in the resistant cultivar (Fig 7B). Similar expression patterns were observed for two more creeping bentgrass GLP genes that did not group with known defense-associated GLPs in phylogenetic analysis (S3 Fig).

**Fig 7.**
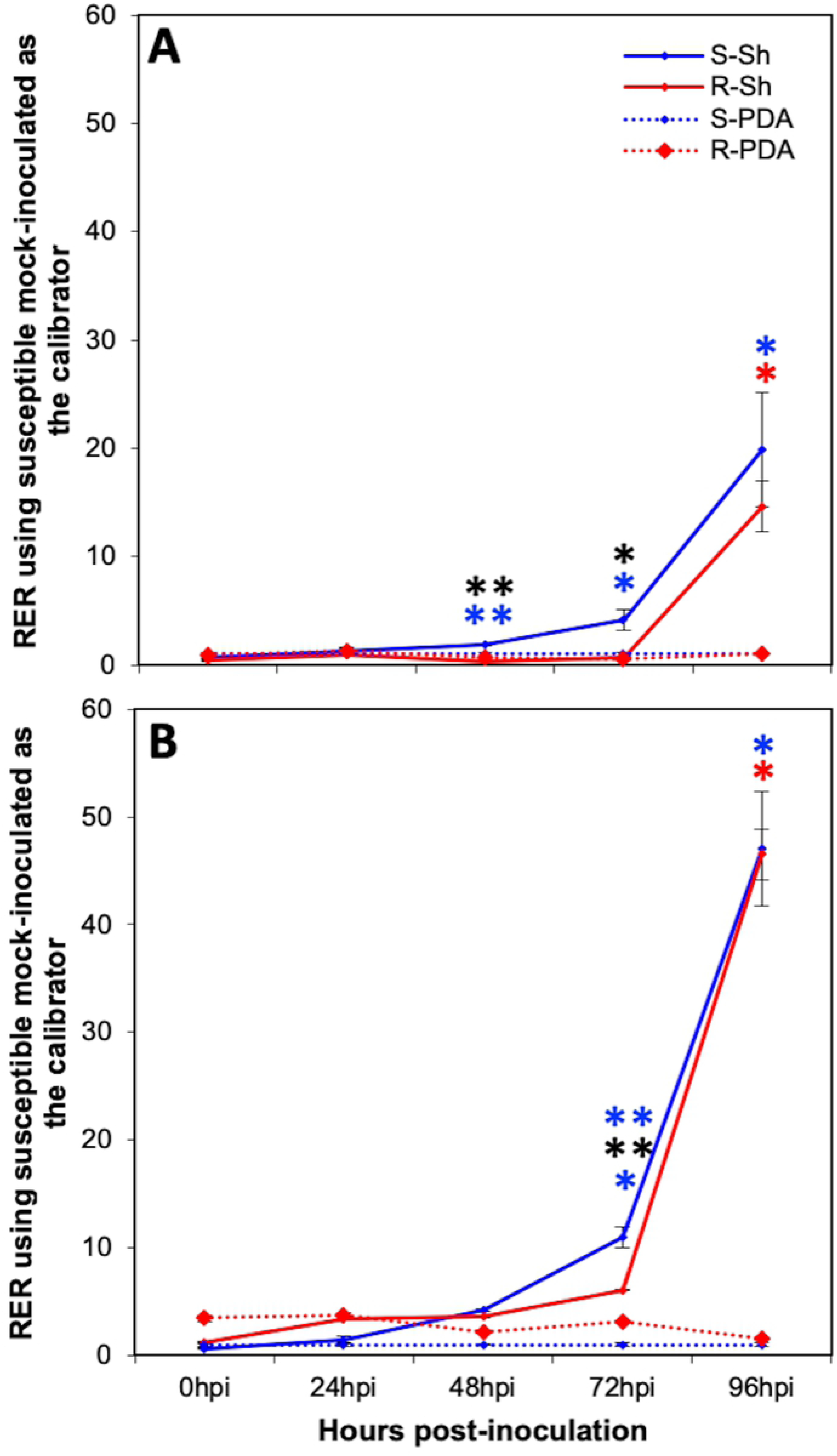
Time-course expression of oxalate oxidase and another germin-like protein gene in resistant and susceptible creeping bentgrass cultivars. **A)** Time-course expression of the creeping bentgrass oxalate gene AST_798. **B)** Time-course expression for the creeping bentgrass GLP GER4 subfamily gene AST_854. Relative expression ratios shown were normalized the constitutively expressed creeping bentgrass ACT7 reference gene. Similar results were obtained with the creeping bentgrass GAPDH gene. In the legend for both **A** and **B**, ‘S’ and ‘R’ represent the ‘susceptible’ and ‘resistant’ cultivar, respectively; ‘Sh’ and ‘PDA’ indicate ‘*Clarireedia* sp.-inoculated’ and ‘PDA mock-inoculated’ samples, respectively. Orthogonal contrasts were used to compare relative expression ratios between specific treatment pairs at the α=0.05 level. Black asterisks indicate a significant difference between relative expression ratios for the *Clarireedia* sp.- inoculated resistant and susceptible cultivar; blue asterisks indicate a significant difference the *Clarireedia* sp. and mock-inoculated samples for the susceptible cultivar; red asterisks indicate a significant difference between the *Clarireedia* sp. and mock-inoculated control for the resistant cultivar. For significance codes, *=P≤0.05; **=P≤0.01; ***=P≤0.001. All error bars represent ± one standard error of the mean.

## Discussion

A primary objective of the present research was to identify a host species for use as a model system to study *Clarireedia* sp. pathogenesis and host resistance. The lack of a significant species by isolate interaction in this research supports previous findings in turfgrass hosts that indicate a lack of race specificity in *Clarireedia* sp./host interactions [10]. Therefore a few model host cultivars or accessions with high and low resistance to dollar spot disease could be used to dissect molecular and physiological aspects of host resistance to this pathogen. We detected differences in disease severity and symptom development between species, indicating that not all are equally suited as a model system for *Clarireedia* sp. and that *A. thaliana* is not a suitable model host for *Clarireedia* sp. The correlation between symptom severity and endogenous oxalate was an unexpected finding. Model hosts with varying levels of endogenous oxalate content may be useful further studies of *Clarireedia* sp./host interactions and how they are affected by endogenous host oxalate levels. Soybean cultivars [40] and spinach varieties [41] differ in oxalate content in the field suggesting natural variation in oxalate levels is common in plant species and should be readily identifiable in a chosen model host system. We are unaware of previous reports that plant endogenous oxalate levels influence resistance to fungal pathogens; however, oxalate is known to influence resistance to herbivory by some insects [42, 43].

To our knowledge, a correlation between host endogenous oxalate content and susceptibility to *Clarireedia* sp. has not previously been demonstrated for creeping bentgrass or any other turfgrass species. Plants produce and use oxalate for a variety of purposes, including defense, pH regulation, osmoregulation, and calcium homeostasis [44] Thus, oxalate content can vary widely between cultivars and environments. Correspondingly, resistance to dollar spot is inherited quantitatively and largely dependent upon environmental conditions [45, 46, 47, 10]. It is possible that oxalate content contributes to dollar spot resistance and this is a hypothesis that should be assessed in cultivars with varying levels of resistance to dollar spot disease.

In addition to genetic differences in creeping bentgrass cultivar oxalate content, management and environmental factors could also be used to alter this host characteristic. It has been reported that fertilizers high in ammonium content and low in nitrate are most effective at suppressing dollar spot in field trials [48]. This is of note because specific forms of nitrogen, in particular nitrate, can affect plant oxalate levels [44]. Promotion or reduction of turfgrass oxalate content could occur in response to different fertilization regimes and suggests a novel means by which nutrient management could be used to manage dollar spot. Further studies in the both the field and lab are needed to better understand the relationship between creeping bentgrass cultivar oxalate content and dollar spot resistance. Additionally, studies that investigate the effects of cultural practices, such as fertilization, on both dollar spot resistance and plant oxalate content will be useful in identifying best management practices for dollar spot suppression. Due to the correlative nature of these studies, it is also possible that it is not the oxalate content specifically that determines the outcome of the host/pathogen interaction but related factors such as foliar pH or pH within the plant.

Germin-like proteins are considered an important part of grass basal defense mechanisms [10, 49]. Consequently, the slow increase in expression of oxalate oxidase and other GLP genes in this study was unexpected. Orshinsky et al. [29] also found strong induction of GLP genes in creeping bentgrass 96 hpi inoculation with *Clarireedia* sp. Interestingly, expression of these genes in creeping bentgrass parallels the timing of symptom appearance and increased more rapidly in the susceptible cultivar compared to the resistant cultivar in this study. One possible reason for the delayed expression of host oxalate oxidase genes is that they are activated in the plant as *Clarireedia* sp. transitions from biotrophy to necrotrophy following early stages of pathogenesis. This hypothesis corresponds with our microscopic examinations of the *Clarireedia* sp. infection process, which demonstrated hyphal growth within host tissue prior to cell death. Alternatively, oxalate oxidase and other GLP genes may be expressed by creeping bentgrass only after detection of pathogen-produced OA. *Brassica napus* GLP genes are induced as early as 6 hpi with *S. sclerotiorum* but it is unclear if expression is related to detection of OA or other signs of pathogen infection [51]. Transient induction of GLP gene expression could have occurred at time-points before the 24 h initial sample collection in this study. Other components of experimental design such as age of the host or tissue collection timing post inoculation could have interfered with our ability to detect changes in gene expression at early time points. Functional studies with plants overexpressing or lacking oxalate oxidase and other GLP genes will help to better identify the importance of these genes in host resistance to *Clarireedia* sp.

*Sclerotinia sclerotiorum* isolates deficient in OA production are hypovirulent or non-pathogenic [39, 52]. Similarly, hypovirulent isolates of the chestnut blight fungus, *Cryphonectria parasitica*, produce lower amounts of OA *in vitro* than their virulent counterparts [53]. In this research, we found that a *Clarireedia* sp. isolate deficient in OA production had decreased aggressiveness on creeping bentgrass but not on *B. distachyon* (Fig 6), indicating that OA contributes to aggressiveness but is not a pathogenicity determinant for *Clarireedia* sp. and that *Clarireedia* sp. with reduced OA production may infect more efficiently on hosts with higher endogenous oxalate content. Similarly, the symptoms observed on *A. thaliana* were visibly distinct from those observed on monocots and no increase in oxalate content was found in symptomatic *A. thaliana* leaves, suggesting that other pathogenesis mechanisms, such as cell well degrading enzymes, are likely responsible for infection of this host.

Based on the present research, both host endogenous oxalate content and pathogen-produced OA affect interactions between *Clarireedia* sp. and its hosts. Results indicating that creeping bentgrass cultivars with resistance to dollar spot have lower oxalate levels than susceptible cultivars is the first indication that this physiological host characteristic is correlated with dollar spot resistance. Further studies on the relationship between oxalate content and resistance to dollar spot are needed, but this has potential as a quantifiable trait for selection of resistant clones in creeping bentgrass breeding programs. A limitation to our studies was the lack of genetic resources for *Clarireedia* sp. and one of its natural hosts, creeping bentgrass; however, we were able to identify potential model systems for study of *Clarireedia* sp.*/*host interactions in this research. In the future, use of a model host system and development of functional genetic resources for *Clarireedia* sp. will enable elucidation of the importance of host oxalate and pathogen-produced OA for pathogenesis of this fungus.

## Acknowledgements

We would like to thank the staff and students at the O.J. Noer Turfgrass Research and Education Facility for their efforts in this research. Thank you to the University of Wisconsin-Madison and Department of Plant Pathology for the support and oversight.

## Supporting Information

**S1 Table.** Plants used in this research.

**S2 Table.** Fungal isolates used in this research.

**S3 Table.** Symptom severity rating scale used for *Clarireedia* spp. infection of creeping bentgrass and models hosts.

**S4 Table.** Primers used for RT-qPCR expression analysis of creeping bentgrass GLP genes.

**S1 Figure. Methods used for inoculation of various hosts with *Clarireedia* spp. A)** Inoculation of *Arabidopsis thaliana* leaves with single agar plugs placed mycelia-side down on leaf surfaces. Inoculated plants were covered with a humidity dome to maintain high relative humidity. **B)** *Brachypodium distachyon* plants were inoculated by placing an agar plug colonized with *Clarireedia sp.* mycelium-side down against foliar material and wrapping with parafilm. **C)** Pots of creeping bentgrass were inoculated by collecting leaf blades in the center of the pot into a bundle and placing a single agar plug against the foliar tissue, then wrapping with parafilm. **D)** The most recent fully expanded leaf of barley, rice, and wheat plant were inoculated by placing a single agar plug colonized with *Clarireedia sp.* mycelia side down on the adaxial side of the leaf, then wrapping with parafilm. Mock-inoculations were performed similarly for all species but *Clarireedia sp.* colonized plugs were replaced with fresh PDA plugs.

**S2 Figure. Oxalic acid production by *Clarireedia sp***. **isolates**. Oxalic acid production was measured *in vitro* using bromophenol blue amended medium, which changes from purple to yellow in color as a result of medium acidification. The area of color change was measured by averaging two perpendicular contrasts across the yellow areas. When areas of color change were diffuse, all were measured and the average measurements were combined. Columns represent the pooled means from three experimental repetitions with three replicates per isolate in each repetition (n=9). Error bars represent ± one standard error of the mean. A1623 (red), ML75 (green), and HE10G14 (blue) were selected as isolates with high, moderate, and low oxalic acid production capacities, respectively.

**S3 Figure. Selection of candidate oxalate oxidase and germin-like protein genes. A)** CRB (AST) and representative barley (HvGer) and rice (Os) GLP amino acid sequences from rice and barley were aligned using ClustalW. JalView was used to color sequence alignment according to percent identity. Key features of GLP genes are annotated with green boxes and black text. Blue arrowheads mark the start and end of the GLP cupin domain. ‘AST’ nucleotide sequences were originally identified by Orshinsky et al. 2012. **B)** Maximum parsimony phylogenetic tree (100 bootstraps with sequence order re-arranging over 10 data sets) of CRB and select grass GLP-genes. The percentage of replicate trees in which the associated taxa clustered together in the bootstrap test are shown next to the branches. Rice and barley defense-associated GLP genes are indicated by the green box. Oxalate oxidase-type CRB GLP-genes are indicated by the orange box. RT-qPCR analysis of expression was performed for creeping bentgrass genes marked by an asterisk. **C)** Time-course expression analysis for creeping bentgrass GLP-genes AST_590 and AST_608. Both genes were significantly up-regulated in the *Clarireedia sp*.-inoculated susceptible cultivar relative to the *Clarireedia sp*.-inoculated resistant cultivar and mock-inoculated susceptible cultivar at 72 hpi (*P*<0.05). At 96 hpi, both genes were upregulated in the *Clarireedia sp*.- inoculated susceptible and resistant cultivars relative to the mock-inoculated controls for each cultivar (*P*<0.05). There was no difference in expression of either gene between the *Clarireedia sp*.-inoculated susceptible and resistant cultivars at 96 hpi (*P*>0.10).

